# GASPS: A Multi-Omics Framework for Defining Genomic Aberration-Driven Signatures and Predicting Patient Outcomes in Lung Cancer

**DOI:** 10.1101/2025.08.21.671519

**Authors:** Zijian Zhang, Xiang Wang, Chenyang Li, Robert Taylor Ripley, Jia Wu, Jianjun Zhang, Christopher I Amos, Chao Cheng

## Abstract

Lung cancer is the most common cause of cancer-related death worldwide. Recent advancements in targeted therapies and immunotherapies have achieved remarkable success. However, patient responses to treatments with lung cancer vary substantially. The mutation status of driver genes can direct personalized treatment, but their prognostic value and treatment efficacy are limited. In this study, we developed a statistical framework named Genomic Aberration-Derived Signature for Patient Stratification (GASPS) to characterize the transcriptomic deregulation of driver genomic aberrations and stratify patients. By applying GASPS to The Cancer Genome Atlas Lung Adenocarcinoma (TCGA-LUAD) data, we developed gene signatures for 38 driver genomic aberrations, including gene mutations, amplifications, and deletions. These signatures were applied to independent lung cancer transcriptomic datasets containing a total of 2,226 patient samples. Our results indicated that these driver gene signatures are much more prognostic than their corresponding genomic mutations. Interestingly, the two EGFR-related signatures characterizing EGFR mutation and amplification, respectively, exhibited contrasting associations with prognosis, treatment response, and immune infiltration in the tumor microenvironment. Moreover, the STK11 mutation signature, rather than the mutation status, was found to be predictive of the response and long-term benefit of patients treated with immune checkpoint blockade therapy in lung cancer. This framework is readily applicable to most cancer types using existing data to improve prognostic risk assessment and treatment efficacy by guiding personalized therapies.

## Introduction

Lung cancer is the leading cause of cancer-related deaths in the U.S. and worldwide^1^. In 2024, the American Cancer Society estimated 234,580 new cases and 125,070 deaths from lung cancer in the United States^2^. Due to the majority of patients being diagnosed at an advanced stage, lung cancer is recognized as an aggressive disease with a poor prognosis^3^. In recent years, numerous new treatment options have been developed and many more are undergoing clinical trials for lung cancer, particularly targeted therapies and immune checkpoint blockade therapies (ICBT)^4,5^. Three generations of tyrosine kinase inhibitors targeting EGFR have been developed to treat non-small cell lung cancer (NSCLC) patients harboring mutations that lead to over-activation of the EGFR gene^6^. Additionally, ICBTs, such as anti-PD-1/PD-L1 treatments, have been widely implemented and have significantly improved the survival outcomes of patients with NSCLC^7^. These therapies are utilized as monotherapy or in combination with other treatments (e.g., chemotherapy), in neoadjuvant or adjuvant settings, based on the stage and clinicopathological characteristics of the patients^8^.

Known as a heterogeneous disease, lung cancer patients exhibit substantial variation in prognosis and response to treatment^9^. Therefore, effective biomarkers are essential for identifying patients at high prognostic risk and enhancing treatment outcomes through precision cancer therapy. The intertumoral heterogeneity observed among lung cancer patients can largely be attributed to genomic variations, including somatic mutations and copy number alterations in their genomes^10^. The initiation and progression of lung cancer in each patient is driven by a distinct set of driver genomic aberrations, which, in turn, influence patient prognosis and treatment response. By identifying driver mutations in actionable genes, personalized treatment using targeted therapies can be implemented. This approach has led to the accelerated FDA approval of Vitrakvi (larotrectinib) for treating solid tumors harboring NTRK gene fusions^11^ and Keytruda (pembrolizumab) for treating any solid tumors with microsatellite instability (MSI), regardless of the tissue of origin^12^.

Despite some successful examples, the reliability of gene mutations in guiding the selection of targeted treatments remains uncertain^13,14^. For instance, in breast cancer, approximately 30-40% of patients harbor mutations in the PIK3CA gene. Although preclinical studies have shown that PIK3CA-mutated breast cancer cells are more responsive to PI3K inhibitors^15^, the reported predictive value of PIK3CA mutations in clinical settings remains contradictory. While some studies have demonstrated their predictive value for pan-PI3K inhibitors^16^, others have found no significant predictive benefit^17–19^. Essentially, tumor development is driven by the aberrant regulation of specific oncogenic or tumor-suppressive pathways, which can be deregulated by various mechanisms at both genomic and epigenomic levels. Although driver genomic aberrations, such as gene mutations, amplifications, deletions, and translocations, are the most likely mechanisms behind pathway deregulation during tumor initiation and progression, they are not the only ones. For example, somatic mutations in the TP53 gene have been observed in more than 50% of tumors. However, inactivation of the downstream p53 pathway can also occur through alternative mechanisms, including loss of TP53, hypermethylation of the TP53 promoter, and genomic alterations in other pathway-associated genes^20^. Therefore, relying exclusively on driver genomic aberrations to predict patient treatment outcomes may not provide sufficient accuracy for guiding personalized cancer treatment.

In this study, we developed a novel statistical framework called Genomic Aberration-Derived Signature for Patient Stratification (GASPS) to integrate cancer genomic and transcriptomic data for improved patient stratification. GASPS defines gene signatures by simultaneously modeling the effects of genomic aberrations on gene expression changes in tumor samples. For a specific cancer type, it identifies gene signatures for each commonly observed driver genomic aberration, including gene mutations, amplifications, and deletions. These signatures can then be applied to independent cancer gene expression profiles to calculate patient-specific signature scores, collectively providing an overview of oncogenic and tumor-suppressive pathways in each tumor sample. In this study, we applied GASPS to lung adenocarcinoma (LUAD) data, defining a total of 38 signatures based on The Cancer Genome Atlas (TCGA) data. We demonstrated that these signatures served as reliable biomarkers for predicting patient prognosis and response to targeted or immunotherapy when validated on multiple independent lung cancer datasets.

## Methods

### Datasets Used in This Study

#### TCGA-LUAD Data

The TCGA LUAD dataset was obtained from the FireBrowse database (http://firebrowse.org), including RNA-seq, somatic mutation, copy number variation (CNV), DNA methylation data, and clinical information. RNA-seq data, normalized by the RSEM method^35^, were available for 515 LUAD patients. Somatic mutation data were provided in Mutation Annotation Format (MAF), detailing functionally relevant mutations such as Frame_Shift, Missense, Nonsense, and Splice_Site variants, while synonymous mutations were excluded. CNV data in Segment (SEG) files contained chromosomal segments with copy number alterations. Amplified and deleted genes were identified based on log_2_-transformed copy numbers. Driver genomic aberrations were determined for cancer-related genes present in at least 10% of LUAD samples, referencing the COSMIC database^36^. DNA methylation data, generated using the Illumina Human Methylation 450K BeadChip platform, provided CpG methylation levels. The median value across CpG sites was used to estimate global methylation per sample.

#### TRACERx Data

The TRACERx project investigates the evolutionary dynamics of NSCLC^37^. Processed RNA-seq, somatic mutation, and clinical data for 421 NSCLC patients were obtained from Zenodo (accession ID 7603386)^22^. The dataset includes primary, metastatic, and lymph node samples; however, only primary tumor samples were used in this study.

#### Gentles Meta-data

The Gentles dataset is a meta-cohort comprising seven NSCLC gene expression datasets with patient survival information^38^. The dataset, containing 1,106 NSCLC samples, was downloaded from the Gene Expression Omnibus (GEO) under accession ID GSE63679^39^.

#### Patil-OAK Data

The Phase 3 OAK trial (NCT02008227) evaluated atezolizumab (anti-PD-L1) versus docetaxel in NSCLC patients^40^. RNA-seq profiles from 699 pretreatment tumor samples were obtained from the European Genome-phenome Archive (EGA) (EGAS00001005013)^23^.

A summary of these datasets was included in the Suppl. Table S1. A more detailed description is provided in the Suppl. Methods and Materials.

### The GASPS framework

GASPS is a statistical framework designed to define gene signatures for driver genomic aberrations in cancer. By analyzing the transcriptomic effects of these aberrations in tumor samples, GASPS enables the calculation of patient-specific signature scores, providing insights into tumor-specific biological pathways. The framework consists of three key components:

#### GASPS-C1: Defining gene signatures for driver genomic aberrations

GASPS constructs gene signatures by modeling the effects of genomic aberrations (e.g., mutations, amplifications, deletions) on gene expression. Using a multivariate linear regression model, informative genes are identified based on significant associations between genomic events and gene expression levels. Genes with significant coefficients are assigned weighted profiles (w⁺ and w⁻), which are trimmed to prevent extreme values and rescaled to a [0,1] range. In this study, we identified a total of 38 driver genomic aberrations based on the TCGA-LUAD somatic mutation and copy number variation data (Suppl. Table S2). These aberrations included 7 mutations, 28 amplifications, and 3 deletions of cancer-related genes that are present in at least 10% of tumor samples from the TCGA-LUAD data.

#### GASPS-C2: Calculating signature scores

Once gene signatures are established, they are applied to independent gene expression datasets to compute sample-specific scores. GASPS employs a modified binding association with sorted expression (BASE) algorithm to quantify pathway activity within tumor samples^41,42^. The method ranks genes by expression level and uses cumulative functions to assess the enrichment of informative genes. The deviation between these distributions is normalized against a null model, generating a final signature score (S = S⁺ - S⁻) that reflects pathway activation. In this study, we applied GASPS to three independent lung cancer transcriptomic data or meta-data including a total of 2226 tumor samples. For each sample, we calculated patient-specific scores for each of the 38 gene signatures to characterize its deregulated pathway.

#### GASPS-C3: Predicting clinical outcomes

Computed signature scores serve as candidate biomarkers for predicting clinical outcomes, including patient prognosis and response to therapy. These signatures can be integrated with clinical variables to improve predictive models, facilitating precision oncology approaches. In this study, we performed systematic analyses to examine the prognostic and predictive values of the 38 gene signatures in lung cancer.

By leveraging genomic and transcriptomic data, GASPS provides a robust framework for patient stratification, advancing personalized cancer treatment strategies. More detailed description of this statistical framework is provided in the Suppl. Methods and Materials

### Clustering analysis of gene signature score

We performed a hierarchical clustering analysis using the signature scores for the 1,106 lung cancer samples included in the Gentles dataset^38^. For each of the 38 driver gene signatures, we calculated the patient-specific scores in all samples. The resultant signature score matrix was used as the input data for clustering analysis to identify patient subgroups. The R function heatmap.2 was used to implement this analysis using the default parameter setting.

#### Deconvolution analysis to infer immune infiltration

The ESTIMATE algorithm was used to estimate the purity of tumor samples based on their gene expression profiles^43^. The TIMER algorithm was applied to deconvolute tumor gene expression data to infer infiltration levels of six major immune cell types, including B cells, CD4 T cells, CD8 T cells, neutrophils, macrophages, and dendritic cells^44^.

### Application of previous gene signatures

We applied several previously published gene signatures to characterize tumor samples based on their gene expression data, including cell proliferation^45^, IFN-γ response^45^, and TGF-β response signatures^46^. These signatures have been applied to all TCGA-LUAD tumor samples by Thorsson et al.^47^, and the results from this article were used in our analysis. In addition, the data provided leukocyte and lymphocyte infiltration levels in each TCGA-LUAD sample, which were also included in our analysis.

### Survival analysis

Cox proportional hazard models were applied to analyze the association between signature scores and patient survival outcomes, including overall survival (OS) and progression-free survival (PFS), based on the availability of survival data in each dataset.

Univariable Cox regression models were applied to evaluate the prognostic effect of individual gene signatures using their scores as continuous variables. The multivariable Cox regression models were constructed to further evaluate their prognostic association after adjusting for established clinical variables such as age, gender, and tumor stage. The Kaplan-Meier plots and survival analyses were implemented using the “survival” and “survminer” packages developed for the R platform. Specifically, the “coxph” function was used to construct Cox proportional hazard models. The “survfit” and “ggsurvplot” functions were used to generate Kaplan-Meier survival curves. The “survdiff” function was used to compare the survival differences of two patient subgroups.

### Other statistical analyses

Pearson and Spearman correlation coefficients were used to quantify and estimate the statistical significance of the associations between continuous variables. The Student t-test and the Wilcoxon rank sum test were applied to evaluate the differences in continuous variables (e.g., signature scores) between two patient groups (e.g., responders versus non-responders). When paired data were provided, paired tests were adopted. One-way ANOVA was conducted to compare continuous variables among multiple groups (e.g., tumor stages). All statistical analyses were performed using the R platform (version 4.3.1).

### Data and code availability

All the data sets used in this study are publicly available. The TCGA-LUAD dataset analyzed in this study was obtained from FireBrowse database at http://firebrowse.org.

The TRACERx data analyzed in this study were obtained from Zenodo at accession ID 7603386. The Gentles dataset analyzed in this study was downloaded from the Gene Expression Omnibus (GEO) at GSE63679. The Patil-OAK data analyzed in this study were obtained from the European Genome-phenome Archive (EGA) at EGAS00001005013. The original code for the analysis is publicly available for non-profit academic use as of the publication date. The example codes are deposited in Github: https://github.com/zijianzhang3/GASPS

## Results

### Define gene signatures to systematically model driver genomic aberration

The development of cancer is driven by genomic alterations in oncogenes or tumor suppressors, including gene mutations, amplifications, and deletions. These driver genomic aberrations often occur in tumor samples in a co-occurring or mutually exclusive manner^21^. Within a single tumor, multiple genomic aberrations may coexist, deregulating downstream oncogenic or tumor-suppressive pathways. These deregulated pathways interact and collectively shape the transcriptomic output. The GASPS framework integrates somatic mutation, copy number variation, and transcriptomic data to define driver gene signatures simultaneously. By modeling the transcriptomic effects of all driver genomic aberrations together, GASPS takes into consideration their co-occurrence or mutual exclusivity while adjusting for established clinical factors such as age and tumor stage. These driver gene signatures can be applied to new cancer expression profiles to characterize the deregulated pathways underlying a sample. (Figure 1a) Compared to the original genomic aberrations, the resultant signature scores show stronger correlations with patient clinical outcomes. In this study, we utilized TCGA-LUAD data to define a total of 38 driver gene signatures, including 7 gene mutations, 28 amplifications, and 3 deletions. We then applied these signatures to multiple high-quality NSCLC datasets to evaluate their ability to predict patient prognosis and response to chemotherapy or immunotherapy (Figure 1b).

**Figure 1:**
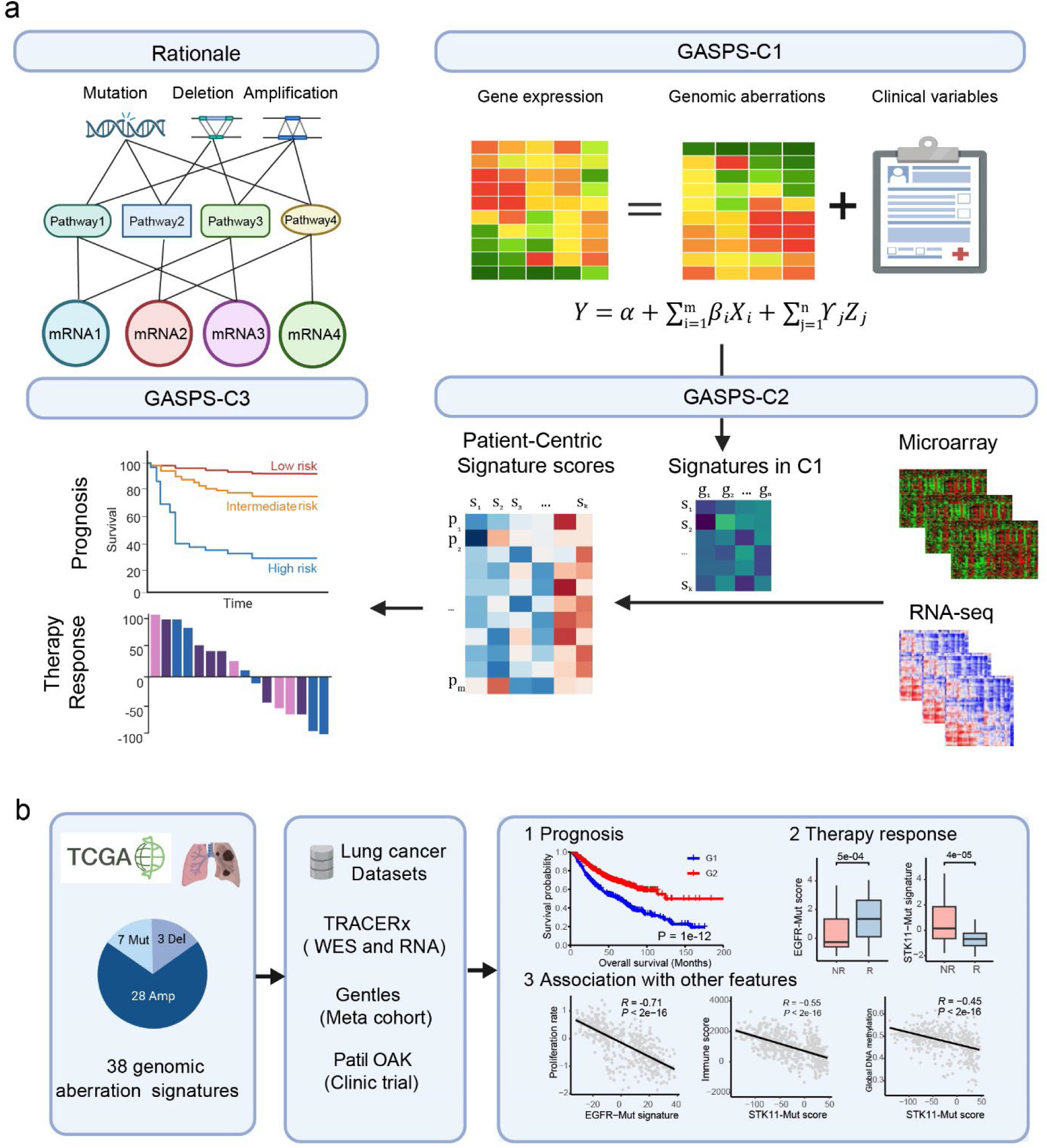
The GASPS framework and its application to lung adenocarcinoma. (a) Overview of the GASPS framework. (b) Application of 38 gene signatures for genomic aberrations to predict patient outcomes in lung adenocarcinoma.

### Validate driver gene signatures using the TRACERx dataset

To validate these signatures, we applied them to an independent RNA-seq dataset generated by the TRAcking Cancer Evolution through Therapy Rx (TRACERx) project. This dataset contains 1,011 multi-regional transcriptomic profiles and whole exome sequencing profiles from a total of 421 patients with NSCLC^22^. Out of the seven genes with defined mutation signatures, six genes—TP53, KRAS, EGFR, STK11, NF1, and SETBP1—are mutated in at least 50 patients (>10%). We expect that the signature scores for a driver mutation would be significantly higher in tumor regions harboring mutations of the corresponding gene compared to those without. Indeed, we validated the effectiveness of all mutation signatures, except for NF1 (Supp. Figure S1). For example, among the 1,011 TRACERx samples, 406 samples have a wild-type *TP53* gene, while 605 have at least one TP53 mutation. Based on the predicted functional impact, the mutated samples were further categorized into 583 with at least one driver mutation in TP53 (Driver-mutation, D) and 22 with only non-driver mutations (Non-Driver mutation, ND). The TP53-Mut signature scores were significantly higher in both the Driver-mutation (*P* = 6e-69) and non-driver mutation (*P* = 6e-07) groups compared to the TP53 wild-type group. Although TP53-Mut signature scores were higher in the Driver-mutation group than in the non-driver mutation group, this difference was not statistically significant (*P* > 0.1). For EGFR, KRAS, and STK11, the corresponding signature scores were significantly higher in the Driver-mutation group compared to both the non-driver mutation and wild-type groups, but no significant difference was observed between the latter two groups.

For SETBP*1*, no driver mutations were identified, but 102 samples had non-driver mutations, which showed significantly higher SETBP1-Mut scores than their wild-type counterparts (*P* = 0.002). These results indicate that the mutation signatures defined based on TCGA-LUAD data can be effectively applied to other datasets to infer gene mutation status and the underlying pathway activities.

### Association of driver gene signatures with patient prognosis

We applied these signatures to the Gentles dataset, a LUAD meta-dataset comprising transcriptomic profiles of 1,086 lung adenocarcinoma samples from seven independent cohorts. For each sample, we calculated a vector of signature scores using the 38 driver gene signatures, transforming the gene expression matrix into a signature score matrix. Hierarchical clustering based on this matrix stratified patients into two distinct groups (Figure 2a), which exhibited significant differences in patient OS (Figure 2b). Next, we assessed the prognostic relevance of individual signatures, identifying 24 signatures that were significantly associated with patient OS, including 13 protective signatures (Hazard Ratio, HR < 1 and *P* < 0.01) and 11 hazardous signatures (HR > 1 and *P* < 0.01) (Figure 2c). For example, the CDKN2A-Del signature score was strongly linked to poor prognosis (*P* = 8e-11), with the high-score group showing significantly shorter OS when dichotomized by the median score (Figure 2d, Suppl. Table S3). Conversely, the EXT1-Amp signature score was associated with favorable prognosis (*P* = 6e-12), where the high-score group displayed significantly longer OS (Figure 2e, Suppl. Table S3).

**Figure 2:**
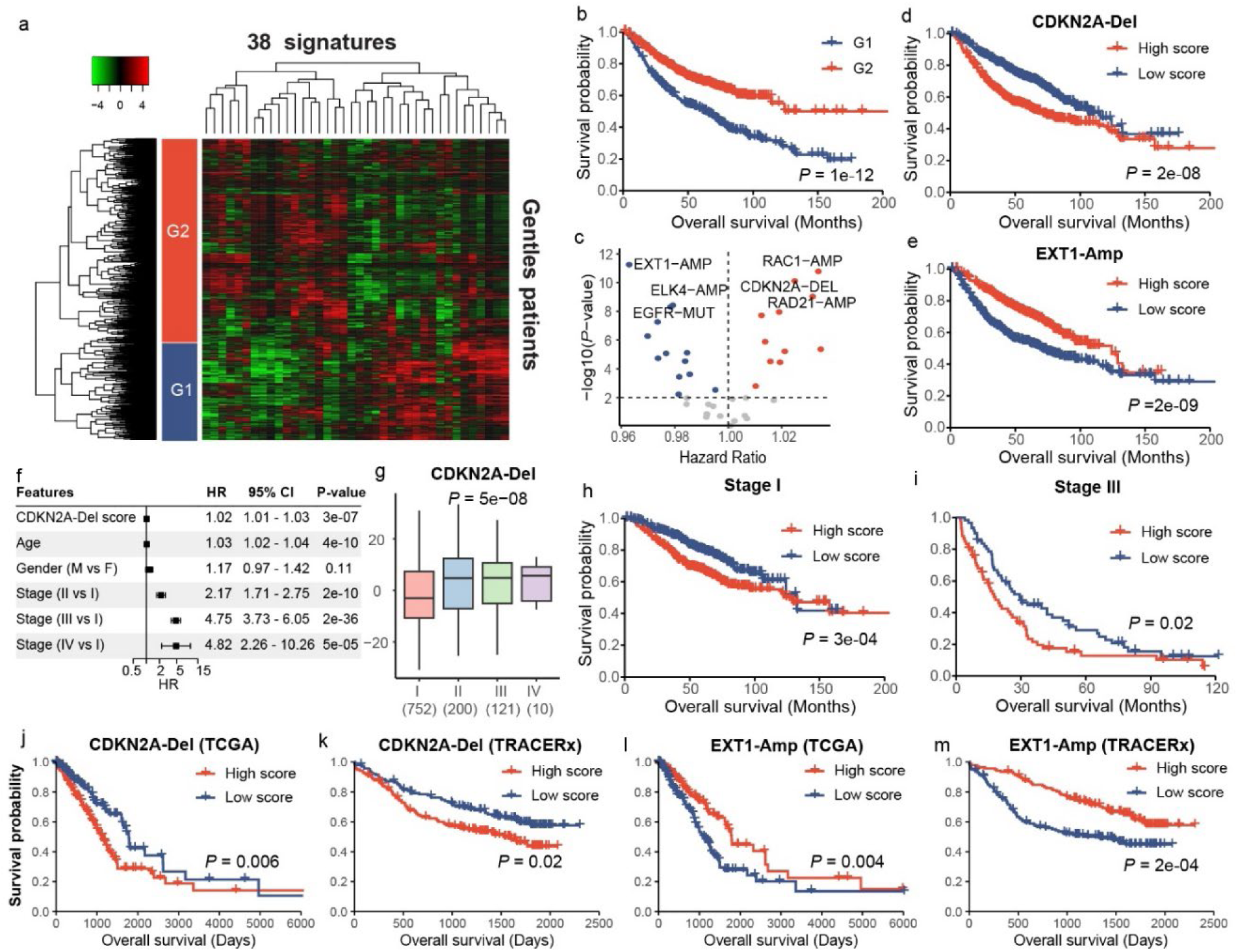
Prognostic value of genomic aberration-derived gene signatures in lung adenocarcinoma. (a) Hierarchical clustering of LUAD samples based on gene signature scores stratified patients into two distinct groups. (b) Kaplan-Meier survival curves exhibiting significant overall survival between the two patient groups. (c) Volcano plot displaying the prognostic association of individual gene signatures. (d-e) Kaplan-Meier survival curves for representative hazardous (CDKN2A-Del) and protective (EXT1-Amp) signatures. Patients with high CDKN2A-Del scores had significantly shorter OS, while high EXT1-Amp scores were associated with improved survival. (f) The CDKN2A-Del signature provides additional prognostic values after adjusting for clinical factors. (g) Boxplot illustrating CDKN2A-Del signature scores across different lung cancer stages, showing increasing scores from early to late stages. (h-i) Kaplan-Meier survival curves for CDKN2A-Del in Stage I and Stage III patients, showing significant prognostic stratification. (j-m) Validation of the CDKN2A-Del and EXT1-Amp signatures in independent LUAD datasets (TCGA and TRACERx), confirming their prognostic significance.

We further evaluated whether these gene signatures provided additional prognostic value after adjusting for established clinical factors through multivariable Cox regression analysis. Most signatures remained significant after adjusting for age, sex, and tumor stage (Suppl. Figure S2), exemplified by the CDKN2A-Del signature (Figure 2f). Stratified analyses within specific tumor stages revealed that CDKN2A-Del signature scores progressively increased from early to late stages, with significant differences observed between stages (Figure 2g). Moreover, this signature could further stratify prognosis within Stage I and Stage III patient subgroups (Figure 2h-i). The prognostic utility of our signatures was validated in independent LUAD datasets, including TCGA-LUAD and TRACERx, where CDKN2A-Del and EXT1-Amp signatures demonstrated consistent prognostic significance (Figure 2j-m).

### Predictive value of driver gene signatures to chemotherapy and immunotherapy

After demonstrating the prognostic value of all signatures, we further evaluated their ability to predict patient responses to chemotherapy and immunotherapy using data from clinical trials. Specifically, we applied our gene signatures to the RNA-seq dataset from the Patil-OAK clinical trial, which includes a docetaxel arm (chemotherapy) and an atezolizumab (anti-PD-L1) arm (immune checkpoint blockade therapy, ICBT) for patients with advanced NSCLC^23^. By comparing signature scores between responders and non-responders in the chemotherapy arm, we identified 10 significant signatures (*P* < 0.01) predictive of patient sensitivity to docetaxel (Figure 3a, Suppl. Table S4). Among these, 5 signatures showed elevated scores in responders. For instance, patients with higher MYC-Amp signature scores were more likely to respond to chemotherapy (Figure 3b). Conversely, 5 signatures exhibited significantly lower scores in responders, such as the RAC1-Amp signature (Figure 3c). Notably, our analysis revealed that the EGFR-Mut signature, but not the EGFR-Amp signature, was associated with chemotherapy sensitivity, which will be discussed in a later section.

**Figure 3:**
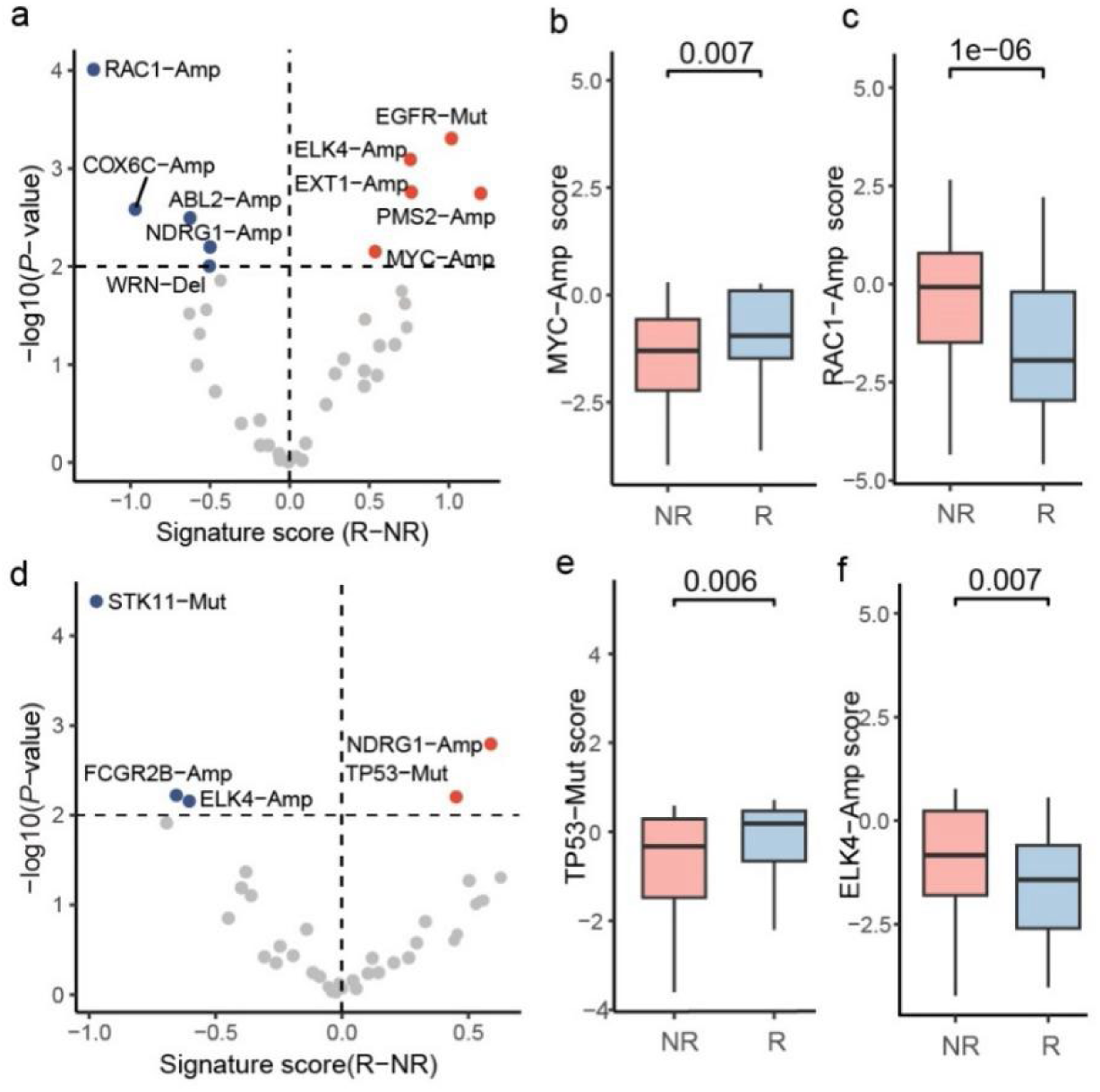
Gene signature scores predict patient response to chemotherapy and immunotherapy. (a) Volcano plot showing gene signatures associated with response to chemotherapy (docetaxel) in the Patil-OAK trial cohort. The X-axis represents the difference in signature scores between responders (R) and non-responders (NR), while the Y-axis indicates the statistical significance (-log10 *P*-value). Red and blue dots denote signatures with significantly higher scores in responders and non-responders, respectively. (b-c) Boxplots showing the distribution of MYC-Amp (b) and RAC1-Amp (c) signature scores between responders and non-responders to chemotherapy. (d) Volcano plot showing gene signatures associated with response to anti-PD-L1 immunotherapy in the same cohort. (e-f) Boxplots depicting the distribution of TP53-Mut (e) and ELK4-Amp (f) signature scores between responders and non-responders to immunotherapy. Statistical significance is indicated above the plots. *R: Responders; NR: Non-responders*.

Similarly, we identified a total of 5 signatures predictive of patient response to anti-PD-L1 therapy, including 2 signatures with elevated scores and 3 signatures with reduced scores in responders compared to non-responders (Figure 3d). For example, patients with a defective p53 pathway, as indicated by higher TP53-Mut signature scores, were more likely to respond to immunotherapy (Figure 3e), while those with higher ELK4-Amp signature scores were more likely to be non-responders (Figure 3f). Notably, the STK11-Mut signature emerged as the most predictive of response, which will also be discussed in detail in a later section. The predictive ability of gene signatures for chemotherapy and immunotherapy was not correlated (*R* = 0.07, Suppl. Figure S3), indicating that distinct sets of signatures are required for response prediction in different treatment modalities.

### Differential prognostic values between EGFR-Mut and EGFR-Amp signatures

Among all gene signatures, the EGFR-Mut and EGFR-Amp signatures were respectively defined based on the mutation and amplification status of the EGFR gene. Our analyses revealed substantial differences in their associations with clinical outcomes and molecular features. First, the EGFR-Mut signature was linked to favorable prognosis, whereas the EGFR-Amp signature was associated with unfavorable prognosis. Specifically, patients with higher EGFR-Mut signature scores demonstrated significantly longer survival times compared to those with lower scores across three independent datasets: Gentles, TRACERx, and TCGA-LUAD (Figure 4a-c). In contrast, the opposite pattern was observed for the EGFR-Amp signature (Figure 4d-f).

**Figure 4:**
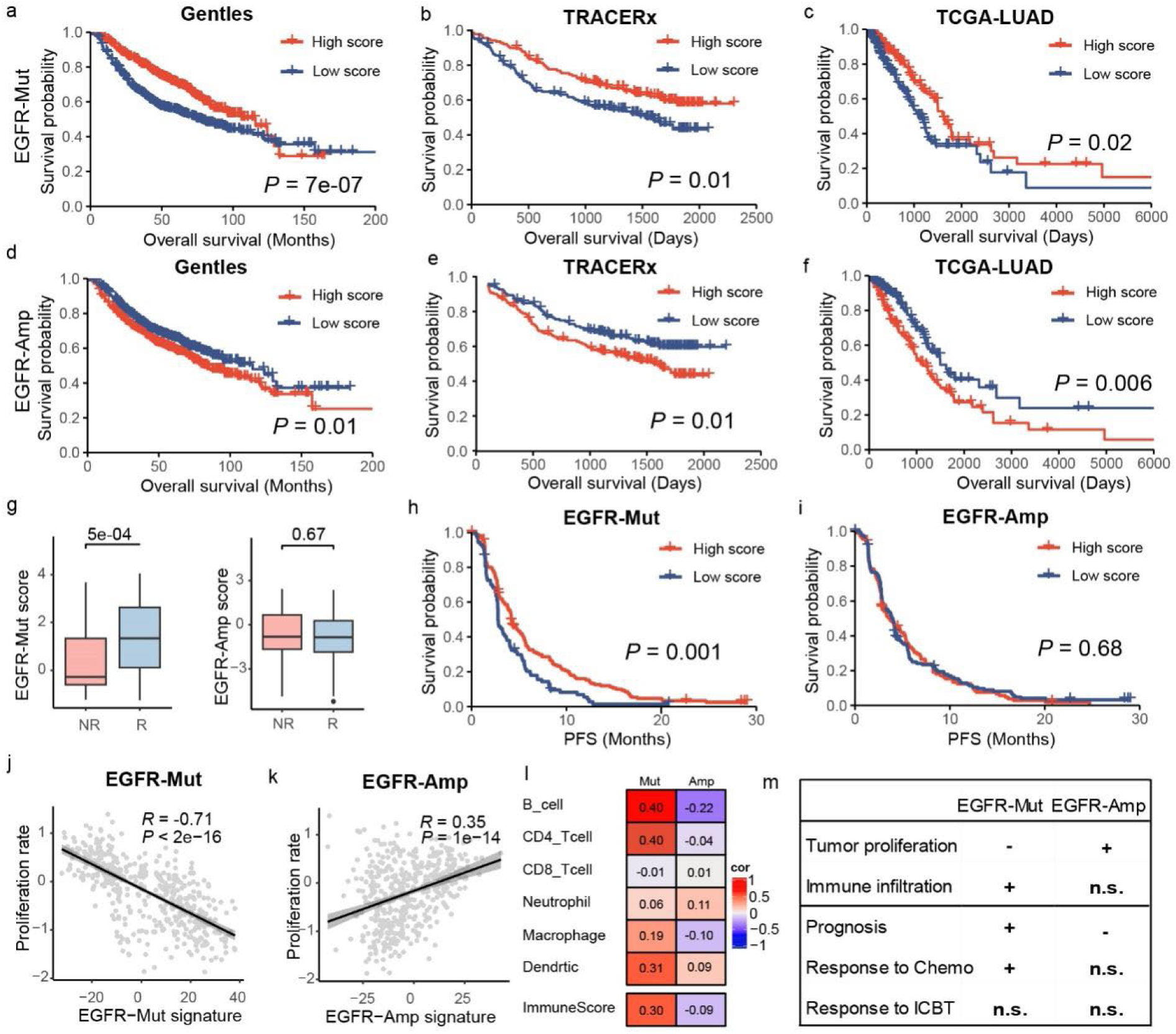
Differential prognostic and predictive value of EGFR-Mut and EGFR-Amp gene signatures in lung cancer. (a-c) Higher EGFR-Mut signature scores were associated with a better prognosis in three independent datasets. (d-f) Higher EGFR-Amp signature scores were associated with a poorer prognosis in the same datasets. (g) EGFR-Mut but not EGFR-Amp signature scores showing significant difference between responders and non-responders to chemotherapy. (h-i) Progression-free survival (PFS) curves for patients treated with chemotherapy, showing that higher EGFR-Mut signature scores are associated with longer PFS, while EGFR-Amp scores show no significant correlation. (j-k) Correlation between EGFR-related signatures and tumor proliferation rates. (l) Heatmap illustrating the correlations of EGFR-Mut and EGFR-Amp signatures with the infiltration level of various immune cell types as well as the overall infiltration level (ImmuneScore). (m) Summary table highlighting differences in tumor proliferation, immune infiltration, prognosis, and treatment response between the two EGFR-related signatures. *R: Responders; NR: Non-responders*.

Second, the EGFR-Mut signature, but not the EGFR-Amp signature, predicted patient sensitivity to chemotherapy in the Patil-OAK dataset. EGFR-Mut signature scores were significantly higher in responders compared to non-responders, a trend not seen for the EGFR-Amp signature (Figure 4g). Notably, patients with higher EGFR-Mut signature scores exhibited longer PFS under chemotherapy, suggesting a sustained benefit from treatment while the EGFR-Amp signature not (Figure 4h-i). However, in the immune checkpoint blockade therapy arm of the same cohort, neither the EGFR-Mut nor the EGFR-Amp signature was correlated with patient response to anti-PD-L1 treatment (Suppl. Figure S4).

To explore the differential associations of these two EGFR-related signatures, we examined their correlations with tumor cell proliferation and immune infiltration in the tumor microenvironment (TME). Interestingly, the EGFR-Mut signature showed a strong negative correlation with tumor proliferation scores (*R* = −0.71, *P* < 0.001), whereas the EGFR-Amp signature exhibited a positive correlation (*R* = 0.35, *P* < 0.001) (Figure 4j-k). Additionally, the EGFR-Mut signature was associated with higher levels of overall immune infiltration (ImmuneScore) and key immune cell populations within the TME (Figure 4l). In contrast, the EGFR-Amp signature showed no significant association with immune infiltration.

Taken together, our findings suggest that the EGFR-Mut and EGFR-Amp signatures have distinct effects on tumor proliferation and immune infiltration, which likely underlie their differing associations with patient prognosis and chemotherapy response (Figure 4m).

### STK11-Mut signature is a predictive biomarker for patient response to ICBT

As described above, the STK11-Mut signature was strongly associated with patient response to ICBT in the Patil-OAK trial. Responders exhibited significantly lower STK11-Mut signature scores than non-responders in the ICBT arm; this difference was absent in the chemotherapy arm (Figure 5a, Supplementary Figure 5). However, response rates did not significantly differ based on STK11 mutation status alone within the ICBT arm (Figure 5b; Fisher’s exact test, P > 0.1). Furthermore, when patients were divided at the median STK11-Mut signature score, the low-score group demonstrated significantly longer PFS than the high-score group, indicating sustained treatment benefits from ICBT (Figure 5c). In contrast, the STK11 mutation itself showed no correlation with PFS in the same cohort (Figure 5d, P > 0.1).

**Figure 5.**
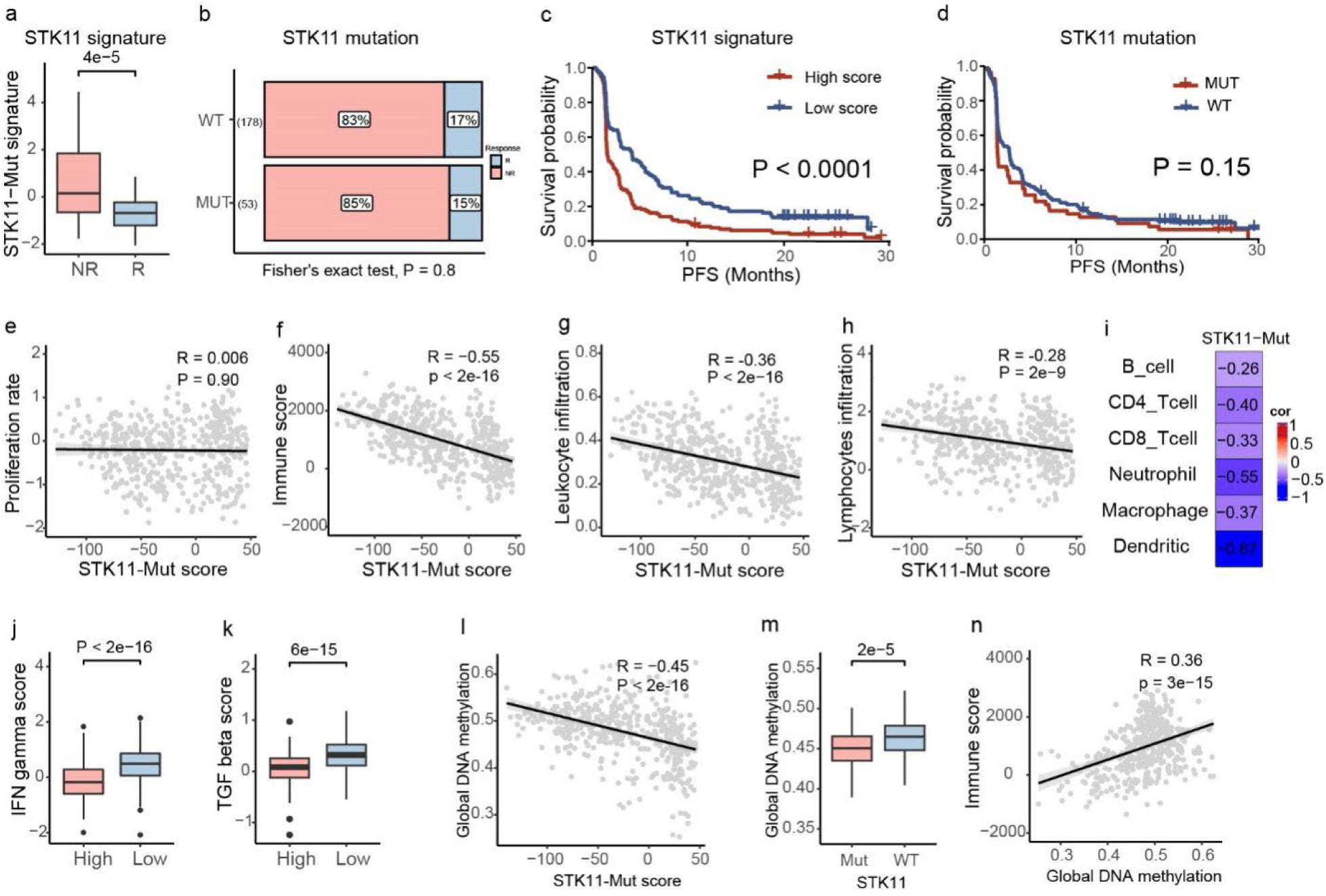
The STK11-Mut signature is predictive of patient response to immune checkpoint blockade therapy (ICBT). (a) STK11-Mut signature scores were significantly lower in responders (R) than in non-responders (NR) to ICBT. (b) STK11 mutation status was not associated with patient response to ICBT. (c-d) Patients with higher STK11-Mut signature scores showed significantly shorter progression-free survival (PFS) when treated by ICBT (c), which was not observed when stratified by the STK11 mutation status (d). (e) Scatter plot showing the lack of correlation between STK11-Mut signature scores and tumor proliferation rate. (f-h) Scatter plots demonstrating significant negative correlations between STK11-Mut signature scores and overall immune score (f), leukocyte infiltration (g), and lymphocyte infiltration (h), suggesting immunosuppressive properties. (i) Heatmap depicting the correlation of STK11-Mut signature scores with different immune cell populations in the tumor microenvironment. (j-k) Boxplots showing significantly higher IFN-gamma (j) and TGF-beta (k) scores in patients with low STK11-Mut signature scores. (l) Scatter plot illustrating a negative correlation between STK11-Mut signature scores and global DNA methylation levels. (m) Boxplot comparing global DNA methylation levels between STK11-mutant and wild-type (WT) samples, showing significantly increased methylation in STK11-mutant cases. (n) Scatter plot showing a positive correlation between global DNA methylation and immune score.

To further investigate the underlying mechanism, we conducted additional analyses. Interestingly, the STK11-Mut signature score was not associated with tumor proliferation (*R* = 0.006) (Figure 5e). However, it exhibited a strong negative correlation with the ImmuneScore (*R* = −0.55), which reflects the overall level of immune infiltration in the TME (Figure 5f). Specifically, the STK11-Mut signature was inversely correlated with the infiltration of both leukocytes (*R* = −0.36) and lymphocytes (*R* = −0.28) (Figure 5g-h). Deconvolution analysis further revealed a negative correlation between this signature and the infiltration of all major immune cell types (Figure 5i). Moreover, patients with higher STK11-Mut scores tended to exhibit lower IFN-γ (Figure 5j) and TGF-β (Figure 5k) responses, suggesting a relatively "cold" immune TME in these samples.

A previous study reported that *STK11* loss in LUAD is associated with global DNA hypomethylation^24^. Using the TCGA-LUAD dataset, we calculated the correlation between all gene signatures and global methylation levels (measured as the median methylation fraction of all CpG sites). The STK11-Mut signature emerged as the most strongly correlated signature (Figure 5l, R = −0.45). Indeed, patients with *STK11* mutations exhibited significantly lower median methylation levels compared to their wild-type counterparts (Figure 5m). Interestingly, global methylation levels were also positively correlated with ImmuneScore (Figure 5n, *R* = 0.36). However, the correlation between STK11-Mut scores and ImmuneScore (*R* = −0.55) was much stronger and remained significant even after adjusting for global methylation levels (Partial Correlation Coefficient = −0.47). Although the global reduction in DNA methylation observed in *STK11*-mutated samples is noteworthy, it does not account for the cold immune environment present in these tumors.

## Discussion

In this study, we developed an integrative framework to model the downstream effects of driver genomic aberrations while accounting for their co-occurrence, mutual exclusivity, and clinical variables. Using TCGA-LUAD data, we defined gene signatures for 38 driver genomic aberrations. These signatures were applied to several lung cancer datasets to transform gene expression profiles into signature score profiles, providing a comprehensive landscape of deregulated pathways in each sample. Compared to the genomic aberrations themselves, these signatures proved more informative in predicting clinical outcomes. Many signatures demonstrated reproducible prognostic value, even when adjusted for established clinical factors. Additionally, some signatures were predictive of patient response to chemotherapy or immunotherapy. Interestingly, despite representing the same gene, we found that the EGFR-Mut and EGFR-Amp signatures differed significantly in their ability to predict clinical outcomes. Finally, we showed that the STK11-Mut signature could serve as a candidate biomarker for predicting patient sensitivity to immunotherapy in NSCLC.

An interesting finding of this study is the differential clinical outcome association between the two *EGFR* signatures. While the EGFR-Mut signature was associated with favorable prognosis under standard care and predictive of patient response to chemotherapy, the EGFR-Amp signature was linked to unfavorable prognosis. These results highlight the distinct downstream effects of these two driver genomic aberrations. Most somatic mutations in the EGFR gene occur at specific hotspots, such as short deletions in exon 19 and the L858R point mutation, which lead to over-activation of the EGFR protein. In contrast, EGFR amplification typically involves broad amplification of chromosomal segments rather than focal amplification of the EGFR gene. Consequently, the EGFR-Amp signature reflects the combined downstream effects of many genes co-amplified with EGFR, rather than EGFR alone. Indeed, the EGFR-Mut and EGFR-Amp signature scores exhibited a negative correlation across all datasets (e.g., *R* = −0.39 in the Gentles dataset). These differences might also account for the inverse correlations between the two signatures and tumor proliferation scores, as well as the association of the EGFR-Mut signature-but not the EGFR-Amp signature-with a cold immune environment in the TME.

A critical discovery from this study is the identification of the STK11-Mut signature as a potential biomarker for predicting patient sensitivity to immune checkpoint blockade therapy (ICBT). Previous studies have reported a link between *STK11* mutations and resistance to immunotherapy^25–27^. STK11 is a tumor suppressor gene, with mutations observed in approximately 17-23% of patients with NSCLC^28–30^. Our results indicated that inactivation of the STK11 gene is associated with patient resistance to anti-PD-L1 treatment, potentially mediated by a cold immune environment in these patients. The association between STK11 mutations and immunotherapy resistance has been noted in prior research^31–33^. An interesting characteristic of STK11 mutations is global DNA hypomethylation, which has been reported in several studies^24,34^. In this study, we demonstrated that global methylation levels of LUAD samples are positively correlated with overall immune infiltration levels in the TME (*R* = 0.36). However, further investigation suggested that global DNA hypomethylation may not be the primary driver of reduced immune infiltration in STK11-mutated samples.

In summary, we developed a robust statistical framework to integrate somatic mutation and copy number variation data with transcriptomic profiles of cancer samples. The resulting gene signatures offer a synergistic representation of deregulated cancer-related pathways associated with a comprehensive list of driver genomic aberrations in a given cancer type. Although demonstrated in LUAD, this framework provides a powerful tool that can be readily applied to other cancer types.

## Author contributions

Z.Z. performed formal analysis, investigation, validation, visualization, and contributed to original drafting and manuscript revision. X.W. conducted formal analysis, investigation, visualization, and contributed to manuscript revision. C.L. contributed to investigation, data curation, and manuscript revision. R.T.R., J.W., J.Z., and C.I.A. contributed to the investigation and critical review of the manuscript. C.C. conceived and supervised the study, developed the methodology, performed formal analysis, contributed to writing the original draft, manuscript revision, and obtained funding. All authors reviewed the manuscript.

## Acknowledgments

This study is supported by the Cancer Prevention Research Institute of Texas (CPRIT) (RR180061) to C.C. and the National Cancer Institute of the National Institutes of Health (1R01CA269764) to C.C. C.C. is a CPRIT Scholar in Cancer Research.

## Declaration of interest

R.T.R. reported grant funding from the National Institutes of Health (R37, R01, U01, U24) and the DeGregorio Family Foundation; clinical trial support from AstraZeneca and Genentech; Vice President of the Board of Directors for the Meso Foundation; and speakers’ bureau for Merck. J.Z. reports research funding from Helius, Johnson and Johnson, Merck, Novartis, Summit, and personal fees from AstraZeneca, Catalyst, GenePlus, Hengrui, Innovent, Johnson and Johnson, Novartis, Oncohost, Takeda and Varian outside the submitted work. The remaining authors declare no conflict of interest.

## Supplemental information titles and legends

### Document Supplementary Methods and Materials

**Supplementary Figure S1-S3**

**Supplementary Table S1-S4**

**Supplementary Figure S1: Driver gene mutation status reflected by genomic aberration signature scores.** (a-f) Boxplots showing signature scores derived from TP53-Mut, EGFR-Mut, KRAS-Mut, STK11-Mut, SETBP1-Mut, and NF1-Mut (from left to right) in the TRACERx dataset, categorized by driver mutations (D), non-driver mutations (ND), and wild-type (WT). Driver mutation scores are significantly higher than wild type except for SETBP1 and NF1. However, non-driver mutation samples show significantly elevated scores compared to wild type in SETBP1. Sample sizes for each group are shown below the boxplots. Statistical significance is denoted by the p-values above each comparison.

**Supplementary Figure S2: The prognosis of genomic aberrations-derived signatures in lung cancer.** (a) Volcano plot showing the association of each gene signatures with prognosis after adjusting for age, gender, and tumor stage using multivariable Cox regression model.

**Supplementary Figure S3: The treatment response prediction performance.** (a) ROC curve showing the predictive performance of MYC-Amp and RAC1-Amp signatures as predictor factors for chemotherapy response. (b) ROC curve exhibiting TP53-Mut and ELK4-Amp signatures in predicting immunotherapy response. (c) Scatter plot displaying no significant correlation of AUC values between chemotherapy response and immune therapy response predictions for each signature.

**Supplementary Figure S4:** The predictive value of EGFR-Mut and EGFR-Amp signatures for immune checkpoint blockade therapy. (a-b) Boxplots showing that no significant differences in EGFR-Amp (a) and EGFR-Mut (b) scores between responders and non-responders in the Patil-OAK immunotherapy arm, respectively. *R: Responders; NR: Non-responders*.

**Supplementary Figure S5:** The predictive value of STK11-Mut signatures for chemotherapy. (a) Boxplots showing that the STK11-Mut signature was not associated with patient response to chemotherapy. (b) Patients with high STK11-Mut signature scores did not show a difference in progression-free survival compared to those with low scores. R: Responders; NR: Non-responders.

**Supplementary Table S1: Datasets used in this study.**

**Supplementary Table S2: Genomic aberrations selected in TCGA-LUAD.**

**Supplementary Table S3: The Cox regression model for signatures in the Gentles dataset.**

**Supplementary Table S4: The difference in signature scores between responders (R) and non-responders (NR) in the Patil-OAK dataset.**

