## Supplementary_Methods_Materials for "GASPS: A Multi-Omics Framework for Defining Genomic Aberration-Driven Signatures and Predicting Patient Outcomes in Lung Cancer"

**Supplemental Methods and Materials**

### Datasets used in this study

*The TCGA-LUAD data*

All TCGA Lung Adenocarcinoma (LUAD) data were generated by the Cancer Genome Atlas (TCGA) project and were downloaded from the FireBrowse database (http://firebrowse.org). The data includes RNA-seq, somatic mutation, copy number variation (CNV), DNA methylation data, and matched clinical information. Level 3 RNA-seq data provide gene expression profiles normalized by the RSEM method for 515LUAD patients[1].

The somatic mutation data and copy number variation (CNV) data were provided as the Mutation Annotation Format (MAF) and segment (SEG) files, respectively. The MAF files include the somatic mutation details for all genes in each sample. Based on these data, we identified mutated genes as those with at least one mutation of the following categories: Frame_Shift_Del, Frame_Shift_Ins, In_Frame_Del, In_Frame_Ins, Missense_Mutation, Nonsense_Mutation, Splice_Site, and Nonstop_Mutation. Synonymous mutations and other mutation categories known to have low functional impact were excluded. The SEG files include chromosome segments significantly deviating from normal copy numbers (n=2) in each sample and provide their inferred copy numbers. These data were used to determine the copy number of genes based on their positions in the genome. Genes with log2 transformed copy numbers greater than $\log_2(3/2)$ and less than $\log_2(1.2/2)$ were identified as amplified and deleted genes, respectively. Based on the gene-level data, we identified driver genomic aberrations associated with cancer genes and presented in at least 10% of LUAD tumor samples. Cancer genes were obtained by referring to the Catalogue Of Somatic Mutations In Cancer (COSMIC) database[2].

TCGA DNA methylation data were generated by using the Illumina Infinium Human Methylation 450K BeadChip platform. The processed data provide the methylation levels (percentage of methylated CpGs) for all measured CpG sites in each sample. The median value of all the CpG sites in a tumor sample was used to represent its global DNA methylation level.

*The TRACEx data*

The TRACERx (TRAcking Cancer Evolution through therapy Rx) project is a longitudinal study investigating the evolutionary dynamics of non-small cell lung cancer (NSCLC)[3]. The TRACEx data provide multiregional RNA and WES sequencing data for tumor samples from 421 patients with non-small cell lung cancer (NSCLC). The processed RNA-seq, somatic mutation, and clinical data for TRACEx were obtained from the Zenodo data repository under accession ID 7603386[4]. The data include primary, metastatic, and lymph node samples. Only data from primary tumor samples were used in our analyses.

*The Gentles meta-data*

The Gentles dataset is a meta-cohort of NSCLC that includes seven datasets with microarray gene expression profiles and matched patient survival information[5]. We downloaded this dataset from the Gene Expression Omnibus (GEO) under accession ID GSE63679[6], which provides gene expression profiles for a total 1,106 NSCLC samples.

*The Patil-OAK data*

The Phase 3 Oak trial (NCT02008227) evaluated the efficacy of atezolizumab, an anti-PD-L1 monoclonal antibody, with comparison to docetaxel in patients with NSCLC[7]. The gene expression profiles for 699 pretreatment tumor samples (each from an individual patient included in the trial) were generated using RNA-seq analysis[8]. The processed RNA-seq data and matched clinical information were downloaded from the European Genome-phenome Archive (EGA) data repository under the accession ID EGAS00001005013.

**The GASPS framework**

The Genomic Aberration-Derived Signature for Patient Stratification (GASPS) is a statistical framework that defines gene signatures for driver genomic aberrations associated with a cancer type. These signatures were developed by examining the transcriptomic effects of genomic aberrations in tumor samples. They can be applied to independent cancer gene expression data to calculate patient-specific signature scores. The resultant signature scores act as proxies for the abnormal molecular pathway alterations, providing insights more correlated with the tumor-specific biological features. The GASPS framework consists of three major components.

*GASPS-C1: Define gene signatures for characterizing driver genomic aberrations.*

GASPS defines gene signatures by jointly modeling the effect of a set of genomic aberrations on gene expression changes. The inputs are matched genomic aberration status at the gene level (including somatic mutations and amplification/deletions of genes) and gene expression profiles for a list of tumor samples. All driver genomic events are considered simultaneously using the following two steps.

1) Identify informative genes. For each gene, we constructed a multivariate linear regression model:

**(Eq. 1)** $$Y = \alpha + \sum_{i=1}^{m} \beta_i X_i + \sum_{j=1}^{n} \gamma_j Z_j,$$

Y is the expression level of the gene; $X_i$ is the indicator function for the genomic event i ($X_i$=1 for samples with the event i, and 0 otherwise); $Z_j$ is the clinical variable j to be adjusted; m and n are the number of genomic events and clinical variable to be considered in the model. By applying the model to the TCGA-LUAD data, we obtained two matrices, $B_{gxm}$ and $P_{gxm}$, containing the coefficients and p-values for all of the g genes to m genomic events, respectively. Informative genes for a genomic event are those with significant p-values in the P matrix.

2) Define gene signatures. For each genomic event, we defined a pair of weighted profiles, denoted as $w^+ = (w_1^+, w_2^+, \ldots, w_g^+)$ and $w^- = (w_1^-, w_2^-, \ldots, w_g^-)$, consisting of all genes. For a gene k with coefficient $b_k$ and p-value $p_k$, we obtained $w_k^+$ and $w_k^-$ by (i) calculating $-I(b_k > 0)\log p_k$ and $-I(b_k < 0)\log p_k$, (ii) trimming at 10 to avoid extreme values, and then (iii) rescaling the values to $[0,1]$.

*GASPS-C2: Calculate signature scores based on gene expression data*

Once defined, these gene signatures ($w^+$and $w^-$) can be used to calculate sample-specific scores to quantify the underlying pathway activity of tumor samples solely based on their gene expression. Specifically, we applied a modified version of a rank-based statistical algorithm named binding association with sorted expression (BASE)[9,10]. First, given the expression profile for a sample, we sorted all genes in the decreasing order of their gene expression, resulting in a ranked gene expression profile $e=(e_1, e_2, \ldots, e_n)$. Second, we examined and quantified the skewed distribution of genes using two cumulative functions, a foreground *f(i)* and a background *b(i).*

**(Eq. 2)** $$f(i) = \frac{\sum_{k=1}^{i} |e_k w_k|}{\sum_{k=1}^{n} |e_k w_k|}, 1 \le i \le n$$

**(Eq. 3)** $$b(i) = \frac{\sum_{k=1}^{i} |e_k (1-w_k)|}{\sum_{k=1}^{n} |e_k (1-w_k)|}, 1 \le i \le n$$

f(i) and b(i) capture the skewed distribution of informative (with high weight) and non-informative (with low weight) genes in a tumor expression profile, respectively. If genes with high weights tend to have higher expression in the profile, f(i) will increase more rapidly with i compared to b(i). Third, we calculated the maximum deviation between f(i) and b(i) and normalized it against a null distribution estimated from permuted data, resulting in a pair of scores, $S^+$ and $S^-$ from $w^+$and $w^-$, respectively. Fourth, the final score for this sample was calculated as $S = S^+ - S^-$.

*GASPS-C3: Use signature scores to predict clinical outcomes*

The resulting signature scores from C2 were used as candidate biomarkers for predicting patient clinical outcomes, including prognosis and patient response to treatment. Informative signatures can be further combined with other features and clinical factors to build integrative prediction models.
