## Supplementary_Figures_Tables for "GASPS: A Multi-Omics Framework for Defining Genomic Aberration-Driven Signatures and Predicting Patient Outcomes in Lung Cancer"

**Supplementary Figures and Tables**

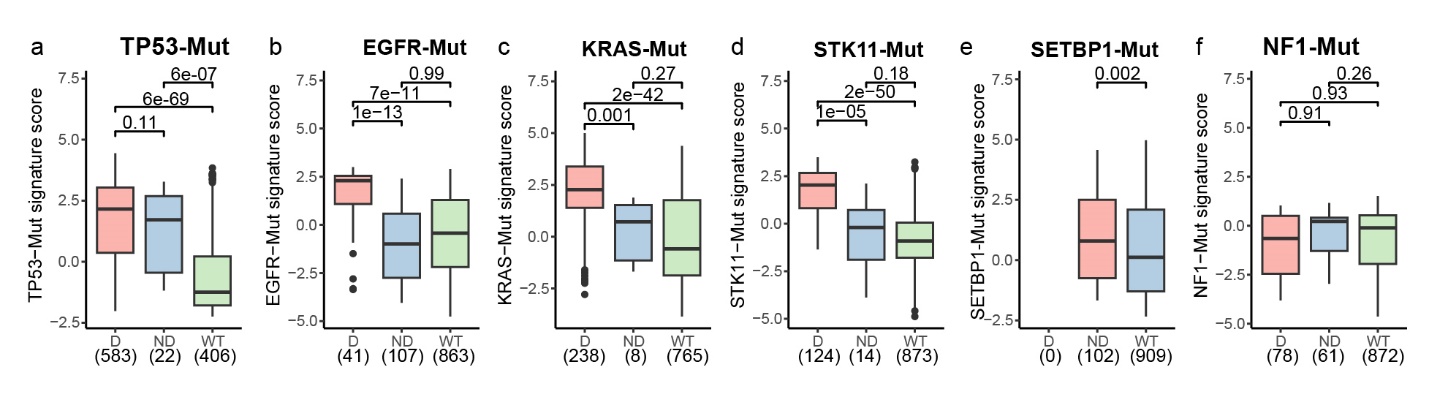

**Supplementary Figure S1: Driver gene mutation status reflected by genomic aberration signature scores.** (a-f) Boxplots showing signature scores derived from TP53-Mut, EGFR-Mut, KRAS-Mut, STK11-Mut, SETBP1-Mut, and NF1-Mut (from left to right) in the TRACERx dataset, categorized by driver mutations (D), non-driver mutations (ND), and wild-type (WT). Driver mutation scores are significantly higher than wild type except for SETBP1 and NF1. However, non-driver mutation samples show significantly elevated scores compared to wild type in SETBP1. Sample sizes for each group are shown below the boxplots. Statistical significance is denoted by the p-values above each comparison.

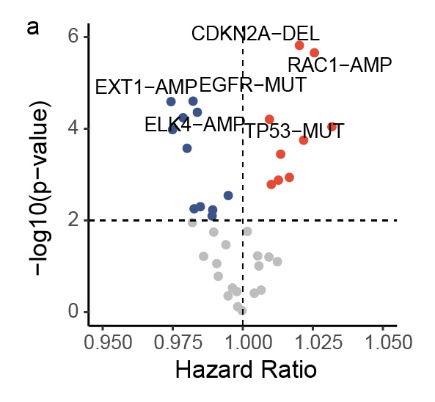

**Supplementary Figure S2: The prognosis of genomic aberrations-derived signatures in lung cancer.** (a) Volcano plot showing the association of each gene signatures with prognosis after adjusting for age, gender, and tumor stage using multivariable Cox regression model.

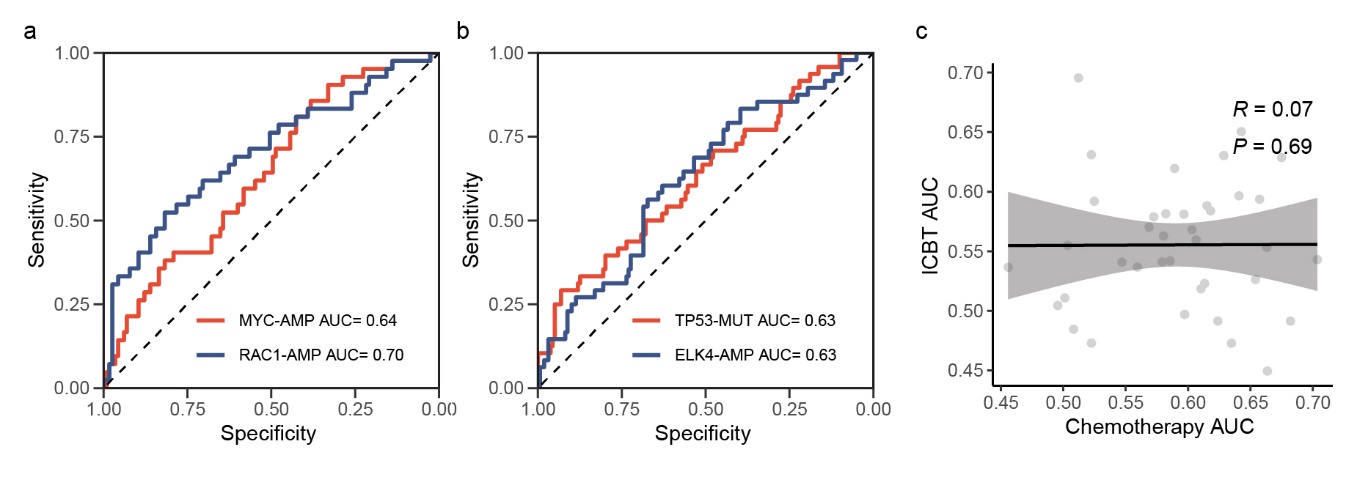

**Supplementary Figure S3:** **The treatment response prediction performance.** (a) ROC curve showing the predictive performance of MYC-Amp and RAC1-Amp signatures as predictor factors for chemotherapy response. (b) ROC curve exhibiting TP53-Mut and ELK4-Amp signatures in predicting immunotherapy response. (c) Scatter plot displaying no significant correlation of AUC values between chemotherapy response and immune therapy response predictions for each signature.

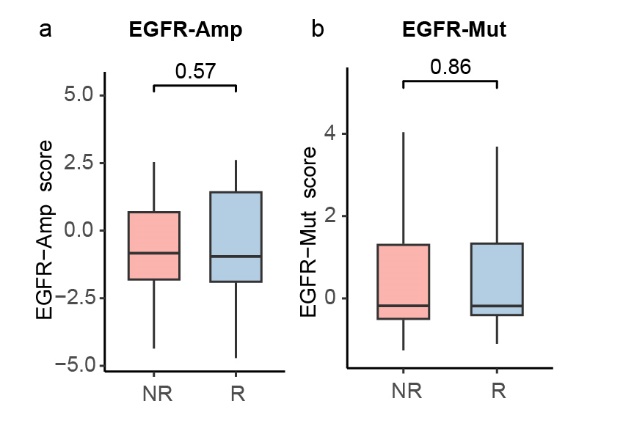

**Supplementary Figure S4:** The predictive value of EGFR-Mut and EGFR-Amp signatures for immune checkpoint blockade therapy. (a-b) Boxplots showing that no significant differences in EGFR-Amp (a) and EGFR-Mut (b) scores between responders and non-responders in the Patil-OAK immunotherapy arm, respectively. *R: Responders; NR: Non-responders.*

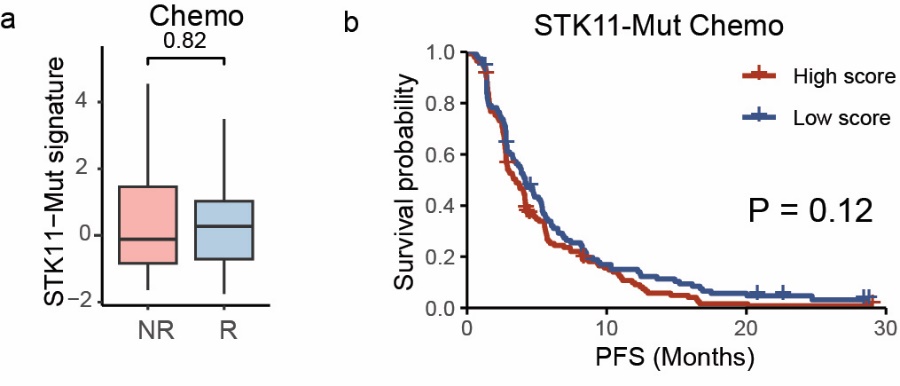

**Supplementary Figure S5:** The predictive value of STK11-Mut signatures for chemotherapy. (a) Boxplots showing that the STK11-Mut signature was not associated with patient response to chemotherapy. (b) Patients with high STK11-Mut signature scores did not show a difference in progression-free survival compared to those with low scores in the chemotherapy arm. *R: Responders; NR: Non-responders.*

**Supplementary Table S1 Datasets used in this study.**

| **Cohort** | **N** | **Samples** | **Treatment** | **Pre-treated biopsy** | **Platform** | **Accession** | **PMID** |
| --- | --- | --- | --- | --- | --- | --- | --- |
| TCGA | 515 | Fresh/frozen | Chemo or target | Yes | NGS | firehose | 25079552 |
| TRACERx | 421 | Fresh/frozen | Naïve | Yes | NGS | zenodo.7603386 | 37046093 |
| Patil-OAK | 699 | FFPE | ICBT or chemo | Yes | NGS | EGAS00001005013 | 35216676 |
| Gentles | 1106 | Fresh/frozen | Chemo or radiation | No | Microarray | GSE67639 | 26286589 |

**Supplementary Table S2 Genomic aberrations selected in TCGA-LUAD.**

| **Signatures** | **Gene** | **Aberration type** |
| --- | --- | --- |
| ABL2-Amp | ABL2 | Amplification |
| ARNT-Amp | ARNT | Amplification |
| CARD11-Amp | CARD11 | Amplification |
| CDKN2A-Del | CDKN2A | Deletion |
| COX6C-Amp | COX6C | Amplification |
| EGFR-Mut | EGFR | Mutation |
| EGFR-Amp | EGFR | Amplification |
| ELK4-Amp | ELK4 | Amplification |
| EXT1-Amp | EXT1 | Amplification |
| FCGR2B-Amp | FCGR2B | Amplification |
| HEY1-Amp | HEY1 | Amplification |
| IKZF1-Amp | IKZF1 | Amplification |
| IL7R-Amp | IL7R | Amplification |
| KRAS-Mut | KRAS | Mutation |
| LIFR-Amp | LIFR | Amplification |
| MDM4-Amp | MDM4 | Amplification |
| MUC1-Amp | MUC1 | Amplification |
| MYC-Amp | MYC | Amplification |
| NCOA2-Amp | NCOA2 | Amplification |
| NDRG1-Amp | NDRG1 | Amplification |
| NF1-Mut | NF1 | Mutation |
| NKX2.1-Amp | NKX2.1 | Amplification |
| NTRK1-Amp | NTRK1 | Amplification |
| PBX1-Amp | PBX1 | Amplification |
| PCM1-Del | PCM1 | Deletion |
| PDE4DIP-Mut | PDE4DIP | Mutation |
| PMS2-Amp | PMS2 | Amplification |
| PTPRC-Amp | PTPRC | Amplification |
| RAC1-Amp | RAC1 | Amplification |
| RAD21-Amp | RAD21 | Amplification |
| SDHC-Amp | SDHC | Amplification |
| SETBP1-Mut | SETBP1 | Mutation |
| STK11-Mut | STK11 | Mutation |
| TERT-Amp | TERT | Amplification |
| TP53-Mut | TP53 | Mutation |
| TPM3-Amp | TPM3 | Amplification |
| TPR-Amp | TPR | Amplification |
| WRN-Del | WRN | Deletion |

**Supplementary Table S3 The Cox regression model for signatures in the Gentles dataset.**

| **Signatures** | **Univariable** | | **Multivariable** | |
| --- | --- | --- | --- | --- |
|  | **Hazard ratio** | **P-value** | **Hazard ratio** | **P-value** |
| ABL2-Amp | 1.14 | 4.E-06 | 1.13 | 9.E-05 |
| ARNT-Amp | 0.87 | 2.E-04 | 0.90 | 8.E-03 |
| CARD11-Amp | 0.94 | 3.E-02 | 0.94 | 6.E-02 |
| CDKN2A-Del | 1.19 | 8.E-11 | 1.15 | 2.E-06 |
| COX6C-Amp | 1.17 | 1.E-08 | 1.12 | 4.E-04 |
| EGFR-Amp | 1.13 | 6.E-06 | 1.10 | 1.E-03 |
| EGFR-Mut | 0.85 | 5.E-09 | 0.87 | 3.E-05 |
| ELK4-Amp | 0.84 | 4.E-09 | 0.88 | 4.E-05 |
| EXT1-Amp | 0.84 | 6.E-12 | 0.89 | 3.E-05 |
| FCGR2B-Amp | 0.96 | 2.E-01 | 0.98 | 4.E-01 |
| HEY1-Amp | 0.87 | 5.E-07 | 0.89 | 1.E-04 |
| IKZF1-Amp | 0.90 | 4.E-04 | 0.92 | 5.E-03 |
| IL7R-Amp | 0.89 | 3.E-03 | 0.88 | 3.E-03 |
| KRAS-Mut | 0.96 | 1.E-01 | 0.95 | 9.E-02 |
| LIFR-Amp | 1.11 | 1.E-02 | 1.12 | 2.E-02 |
| MDM4-Amp | 1.03 | 4.E-01 | 1.00 | 9.E-01 |
| MUC1-Amp | 0.87 | 8.E-06 | 0.91 | 6.E-03 |
| MYC-Amp | 0.89 | 8.E-06 | 0.90 | 3.E-04 |
| NCOA2-Amp | 1.09 | 1.E-02 | 1.07 | 6.E-02 |
| NDRG1-Amp | 1.16 | 3.E-05 | 1.13 | 1.E-03 |
| NF1-Mut | 0.88 | 3.E-05 | 0.92 | 2.E-02 |
| NKX2.1-Amp | 0.92 | 4.E-02 | 0.91 | 3.E-02 |
| NTRK1-Amp | 1.07 | 2.E-02 | 1.05 | 8.E-02 |
| PBX1-Amp | 1.00 | 9.E-01 | 0.99 | 8.E-01 |
| PCM1-Del | 0.92 | 3.E-02 | 0.96 | 3.E-01 |
| PDE4DIP-Mut | 1.11 | 2.E-03 | 1.06 | 1.E-01 |
| PMS2-Amp | 0.88 | 2.E-05 | 0.92 | 6.E-03 |
| PTPRC-Amp | 1.06 | 2.E-01 | 1.04 | 4.E-01 |
| RAC1-Amp | 1.28 | 2.E-11 | 1.21 | 2.E-06 |
| RAD21-Amp | 1.18 | 9.E-10 | 1.12 | 2.E-04 |
| SDHC-Amp | 1.03 | 3.E-01 | 1.03 | 3.E-01 |
| SETBP1-Mut | 0.87 | 6.E-08 | 0.89 | 6.E-05 |
| STK11-Mut | 0.96 | 2.E-01 | 0.97 | 4.E-01 |
| TERT-Amp | 0.93 | 1.E-02 | 0.96 | 2.E-01 |
| TP53-Mut | 1.17 | 2.E-08 | 1.13 | 6.E-05 |
| TPM3-Amp | 1.16 | 1.E-06 | 1.12 | 2.E-03 |
| TPR-Amp | 0.93 | 6.E-03 | 0.93 | 1.E-02 |
| WRN-Del | 1.12 | 3.50E-05 | 1.06 | 6.E-02 |

**Supplementary Table S4 The difference in signature scores between responders (R) and non-responders (NR) in the Patil-OAK dataset.**

| **Signatures** | **Chemotherapy** | | **ICBT** | |
| --- | --- | --- | --- | --- |
|  | **Score diff(R-NR)** | **P-value** | **Score diff(R-NR)** | **P-value** |
| ABL2-Amp | -0.63 | 3.E-03 | 0.10 | 6.E-01 |
| ARNT-Amp | 0.23 | 3.E-01 | 0.21 | 4.E-01 |
| CARD11-Amp | 0.70 | 2.E-02 | 0.06 | 9.E-01 |
| CDKN2A-Del | -0.52 | 3.E-02 | -0.40 | 6.E-02 |
| COX6C-Amp | -0.97 | 3.E-03 | 0.63 | 5.E-02 |
| EGFR-Amp | -0.18 | 7.E-01 | 0.15 | 6.E-01 |
| EGFR-Mut | 1.02 | 5.E-04 | 0.00 | 9.E-01 |
| ELK4-Amp | 0.76 | 8.E-04 | -0.60 | 7.E-03 |
| EXT1-Amp | 0.76 | 2.E-03 | -0.24 | 3.E-01 |
| FCGR2B-Amp | -0.13 | 7.E-01 | -0.66 | 6.E-03 |
| HEY1-Amp | 0.66 | 6.E-02 | -0.02 | 1.E+00 |
| IKZF1-Amp | 0.04 | 9.E-01 | -0.01 | 7.E-01 |
| IL7R-Amp | 0.55 | 1.E-01 | 0.26 | 4.E-01 |
| KRAS-Mut | 0.74 | 4.E-02 | 0.46 | 2.E-01 |
| LIFR-Amp | -0.47 | 2.E-01 | -0.45 | 1.E-01 |
| MDM4-Amp | -0.19 | 4.E-01 | 0.12 | 4.E-01 |
| MUC1-Amp | 0.47 | 2.E-01 | 0.53 | 1.E-01 |
| MYC-Amp | 0.54 | 7.E-03 | -0.38 | 4.E-02 |
| NCOA2-Amp | -0.30 | 4.E-01 | -0.26 | 4.E-01 |
| NDRG1-Amp | -0.50 | 6.E-03 | 0.59 | 2.E-03 |
| NF1-Mut | 0.47 | 1.E-01 | 0.54 | 9.E-02 |
| NKX2.1-Amp | 0.56 | 6.E-02 | 0.56 | 9.E-02 |
| NTRK1-Amp | -0.01 | 1.E+00 | -0.05 | 8.E-01 |
| PBX1-Amp | 0.34 | 9.E-02 | -0.69 | 1.E-02 |
| PCM1-Del | 0.29 | 1.E-01 | -0.14 | 2.E-01 |
| PDE4DIP-Mut | -0.57 | 5.E-02 | 0.33 | 2.E-01 |
| PMS2-Amp | 1.20 | 2.E-03 | 0.30 | 3.E-01 |
| PTPRC-Amp | -0.06 | 9.E-01 | -0.04 | 9.E-01 |
| RAC1-Amp | -1.23 | 1.E-04 | -0.19 | 4.E-01 |
| RAD21-Amp | -0.63 | 3.E-02 | -0.09 | 6.E-01 |
| SDHC-Amp | 0.10 | 6.E-01 | 0.50 | 5.E-02 |
| SETBP1-Mut | 0.47 | 3.E-02 | 0.04 | 7.E-01 |
| STK11-Mut | -0.07 | 8.E-01 | -0.97 | 4.E-05 |
| TERT-Amp | 0.72 | 2.E-02 | -0.36 | 8.E-02 |
| TP53-Mut | -0.43 | 1.E-02 | 0.45 | 6.E-03 |
| TPM3-Amp | -0.58 | 1.E-01 | -0.31 | 4.E-01 |
| TPR-Amp | 0.08 | 9.E-01 | 0.44 | 2.E-01 |
| WRN-Del | -0.50 | 1.E-02 | -0.12 | 6.E-01 |
